# Spatiotemporal Determination of Metabolite Activities in Multiple Corneal Epithelium Barriers on a Chip

**DOI:** 10.1101/2021.01.08.425838

**Authors:** Rodi Abdalkader, Romanas Chaleckis, Craig E. Wheelock, Ken-ichiro Kamei

## Abstract

The corneal epithelial barrier maintains the metabolic activities of the ocular surface by regulating membrane transporters and metabolic enzymes responsible for the homeostasis of the eye as well as the pharmacokinetic behavior of drugs. Despite its importance, no established biomimetic in vitro methods are available to perform the spatiotemporal investigation of corneal metabolism and determine the transportation of endogenous and exogenous molecules. This study introduces multiple corneal epithelium barriers on a chip, namely, Cornea-Chip, which enables the spatiotemporal collection as well as analysis of micro-scaled extracellular metabolites from both the apical and basolateral sides of the barriers. Longitudinal samples collected during 48 h period were analyzed using untargeted liquid chromatography–mass spectrometry metabolomics method, and 104 metabolites were annotated. The shifts in extracellular metabolites and quantitative analysis of the mRNA associated with membrane transporters could allow the investigation of the correlation between the expression of and the secretion and transportation of metabolites across the polarized corneal epithelial barrier. The Cornea-Chip might provide a non-invasive, simple, and effectively informative method to determine pharmacokinetics and pharmacodynamics as well as to discover novel biomarkers for drug toxicological and safety tests as an alternative to animal experiments.

## 1. Introduction

The corneal epithelial barrier is the main structure that determines the pharmacokinetics and pharmacodynamics of drugs into the interior segment of the eye and contributes to the maintenance of homeostasis of the ocular surface by controlling the metabolism and transportation of different molecules from and to the tear pool.^[1,2]^ Nutrients and drugs are actively or passively transported through the corneal epithelial barrier, followed by their intracellular metabolism, modification, and secretion of the corresponding metabolites. Although many metabolic pathways (e.g., oxidative, hydrolytic, reductive, and conjugative pathways)^[3,4]^ and transporters^[5,6]^ in the cornea have been investigated, the spatiotemporal mechanisms of the in situ metabolism and transportation of drugs through the human corneal barrier are barely known, owing to the technical difficulties in the current corneal experimental platforms as well as the lack of analytical tools that non-invasively enable longitudinal metabolome monitoring in small volumes.^[7,8]^ Moreover, while the current in vitro corneal models that use cell-culture inserts (e.g., a transwell system) have been investigated, they only provide a two-dimensional structure of the corneal epithelial cells, which is physiologically different that in the cornea.^[9,10]^ Therefore, in vitro biomimetic models of the cornea need to be established.

These aforementioned requirements can be fulfilled by identifying an alternative in vitro platform that can simulate the structure of the corneal epithelial barrier. Unlike classical models, organ-on-a-chip (OoC) technology is a promising approach for the recreation of miniaturized organs in micro-scaled devices,^[11]^ because it can provide proper threedimensional extracellular environments as well as dynamic flow and mechanical stimuli.^[12–15]^ These advantages also allow the elucidation of the mechanisms of metabolic and transportation activities of the human corneal epithelial barrier to simulate the human corneal epithelial structure and physical stimuli-mediated eye-blinking in vitro. We previously developed an OoC platform with multiple corneal epithelial barriers in a single microfluidic device having apical and basolateral sides under eye blinking-like stimuli, and aqueous humor drainage within the device could characterize both the anatomical structure as well as the biomechanical nature of the corneal epithelial barrier.^[15]^

In addition to the OoC platform, to determine the overall biological function of the corneal epithelial barrier on a chip in a spatiotemporal manner, we need a non-invasive analytical tool that allows the accurate monitoring of enzyme and transporter activities. Many OoC platforms developed to study the metabolism and transport mechanisms use optical or fluorescent assays [e.g., 3-(4,5-dimethylthial-2-yl)-2,5-diphenyltetrazalium bromide (MTT)] and lucifer yellow.^[16–18]^ However, these assays provide very limited information that is restricted to a single target and might cause cellular damage because of the use of additional chemical compounds. In addition to optical and fluorescent assays, untargeted metabolomic analysis based on liquid chromatography–mass spectrometry (LC-MS) is a preferred methodology, since it allows quantification of numerous metabolites in a single measurement.^[19,20]^ Moreover, we have developed an LC-MS-based untargeted metabolomics workflow enabling non-invasive temporal quantification of micro-scaled extracellular metabolites.^[21]^ Thus, the OoC platform in combination with LC-MS untargeted metabolomics might provide deeper insights into the metabolic and transport activities of the improved human corneal epithelial barrier than those provided by conventional cell culture and assay platforms.

Herein, we introduce our OoC platform to reconstruct the human corneal epithelial barrier, namely, Cornea-Chip, for the spatiotemporal collection and analysis of extracellular metabolite secretion and transportation across the barrier. The Cornea-Chip allows the collection of extracellular metabolites separately at the apical and basolateral sides of corneal epithelial cells. Quantitative real time-polymerase chain reaction (RT-PCR) analysis can then be used to evaluate the expression levels of transporters in the corneal epithelial cells. To our knowledge, this is the first study to show that the integration of Cornea-Chip and untargeted metabolomics can allow the assessment of the activities, secretion, and transportation of metabolites across the corneal epithelial barrier.

## 2. Results and Discussion

### 2.1. The construction of Cornea-Chip

To study the process of molecular transport, secretion, and metabolism in the cornea (**Figure 1A**), we constructed the Cornea-Chip by using a multi-well microfluidic device (**Figure 1B**). We confirmed that the corneal cells cultured on a chip formed a thicker 3D structure than that in a cell-culture insert in a shorter period (7 days vs 10–15 days in cellculture inserts). Moreover, the Cornea-Chip allowed the application of eye blinking-like shear stress stimulus on the cultured corneal cells as well as the simulation of the aqueous humor flow.^[15]^ Thus, unlike conventional cell-culture inserts, this chip provides physiologically better conditions for cultured corneal cells retaining their normal functions.

**Figure 1.**
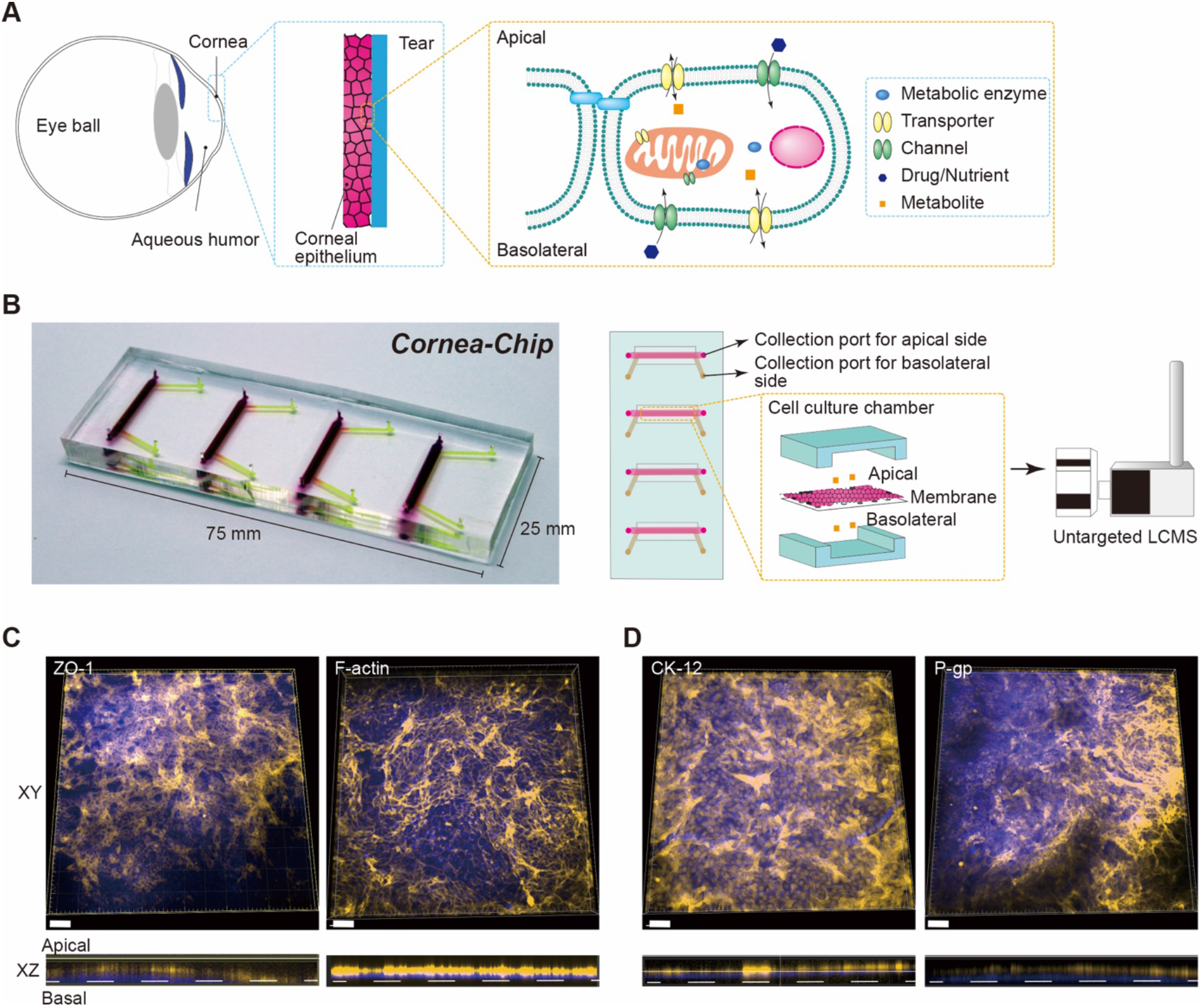
Multiple corneal epithelial barriers on a chip, namely, Cornea-Chip. A) An illustration of the overall structure of the ocular surface, and a scope that shows the role of the corneal barrier in the transportation or metabolism of nutrients and drugs. B) A photograph of Cornea-Chip, and an illustration of its cross-section showing the cell culture chamber from the top to bottom: the upper channel, corneal epithelial cells, porous membrane, and lower channel as well as its applicability for the determination of extracellular metabolites by using untargeted LCMS instrumentation. Each chamber has sample collection ports for apical and basolateral sides. C). Immunofluorescence staining indicating the barrier polarity and intactness (ZO-1 and F-actin). D) Immunofluorescence staining indicating the expression of corneal epithelium maturation marker (CK-12), and the apical ABC-related family of transporter protein (P-gp) expression. XZ dimensions refer to the cross-section scanning images obtained using confocal scanning microscopy. Experimental data are collected from HCE-T cells grown in microfluidic devices for 7 days. Yellow: ZO-1, F-actin, Ck-12, P-gp; blue: DAPI. Scale bar, 100 μm.

The Cornea-Chip consists of four sets of upper and lower channels separated by a clear polyethylene terephthalate (PET) porous membrane (pore size, 0.4 μm; thickness, 10 μm; nominal pore density, 4 × 10^6^ pores cm^−2^). The porous membrane allows the culturing of corneal epithelial cells with distinguished apical and basal sides, as well as exchange of molecules through the pores. The upper and lower layers are made of polydimethylsiloxane (PDMS), which allows high diffusion of gases, thereby providing a suitable environment for cell growth and expansion.^[22]^ The corneal epithelial cells are inoculated in the upper channel having a surface area of 0.23 cm^2^, which is five-times smaller than that of the human cornea (human cornea, 1.32 cm^3^). Molecule exchange across the corneal epithelial barrier formed on the porous membrane is facilitated by the transporting activities of the cultured corneal epithelial cells.

To construct Cornea-Chip, we used human corneal epithelial cells (HCE-T), since these cells express sufficient metabolic enzymes^[23]^ and transmembrane transporters^[24]^ and have long been used for ocular drug development.^[10,25]^ To form the corneal barrier, we seeded the cells in the upper channels and cultured them under static conditions for 7 days. During the 7-day culture on the chip, HCE-T cells proliferated to cover the entire porous membrane (**Figure S1, Supporting Information**). Next, to confirm the intactness and polarity of the barrier, we performed fluorescent immunocytochemistry to analyze the expression of tight junction proteins (zonula occludens protein-1; ZO-1) and F-actin filaments (phalloidin; **Figure 1C**). HCE-T cells cultured on a chip expressed both ZO-1 and F-actin proteins on the apical cellular surface, as in natural corneal epithelial cells.^[26,27]^ Fluorescent immunocytochemistry analysis also revealed that the HCE-T cells cultured on the chip expressed cytokeratin-12 (CK-12, a marker of corneal epithelium maturation) and P-glycoprotein (P-gp, also known as multidrug-resistance protein 1; MDR1, a transmembrane transporter; **Figure 1D** and **Figure S1, Supporting Information**), indicating their ability to form an intact and functional corneal epithelial barrier.

### 2.2. Gene profiling of metabolic transporters on human corneal cells

To investigate metabolic and transporting activities in the Cornea-Chip, we evaluated the mRNA expression of 85 well-known transporters by using a qPCR array in HCE-T cells (**Figure 2**, **Table S2**, **Supporting Information**, and **Methods**). We found the expression of a wide range of transporters (threshold of mRNA expression, 2^−ΔCt^ ≥ 10^−3^) mainly among the genes of the superfamily of solute carrier (SLC)-related transporters, including 14 subgroups (SLC2A1/A2/A3, SLC3A2, SLC5A1, SLC7A5/A6/A7/A8/A11, SLC10A1, SLC16A1/A2/A3, SLC19A1/A2, SLC22A1/A2/A9, SLC25A13, SLC28A1/A3, SLC29A1/A2, SLC31A1, SLC38A2/A5, SLCO2A1/SLCO3A1/SLCO4A1, and SLO1B1/SLO2B1) that mediate the transportation of peptides, monocarboxylic acids, nucleosides, organic anions/cations, glucose, and amino acids.^[28]^ In addition, the ATP-binding cassette (ABC) transporter family, including 6 major subgroups (ABCA2/A5/A9/A12, ABCB1/B4/B6, ABCC1/C3/C4/C5/C16, ABCD1/D4, ABCF1, and ABCG2) responsible for the transportation and efflux of xenobiotics and endogenous molecules, showed remarkable expression.^[29]^ The two subfamilies (VDAC1-C2) of voltage-dependent anion channels (VDACs), Transporter 1, ABC transporter sub-family B [MDR/TAP1-2 and major vault protein (MVP)], which mediate the transportation of xenobiotic toxins, were markedly expressed. Previous studies indicated that some SLC transporters were expressed in the corneal epithelium, mainly SLC22 family members [also known as organic zwitterons/cation transporters (OCTNs) and organic anion transporters (OATs)], owing to their role in the transportation of cationic and anionic drugs.^[2]^ Our results indicated that more SLC subfamilies that are responsible for trafficking of neutral amino acids (SLC38), copper (SLC31), nucleosides (SLC29), monocarboxylate (SLC16), heteromeric amino acids (SLC7), and sugar alcohol (SLC2) were expressed in the corneal cells. In fact, the expression of ABC transporters, mainly ABCC1-C2, ABCG2, and ABCB1, which are known for their activities in the efflux of molecules into the extracellular space, has already been reported in HCE-T cells.^[24,30]^ In addition, we found that the corneal cells expressed ABCC4/C5/C16 (cyclic nucleotide transport) as well as other subgroups such as ABCFD1/D4 (fatty acid transport).^[28]^ Although ABCF1 expression was clearly observed, it might not function as a transporter because of the lack of a putative transmembrane domain.^[31]^

**Figure 2.**
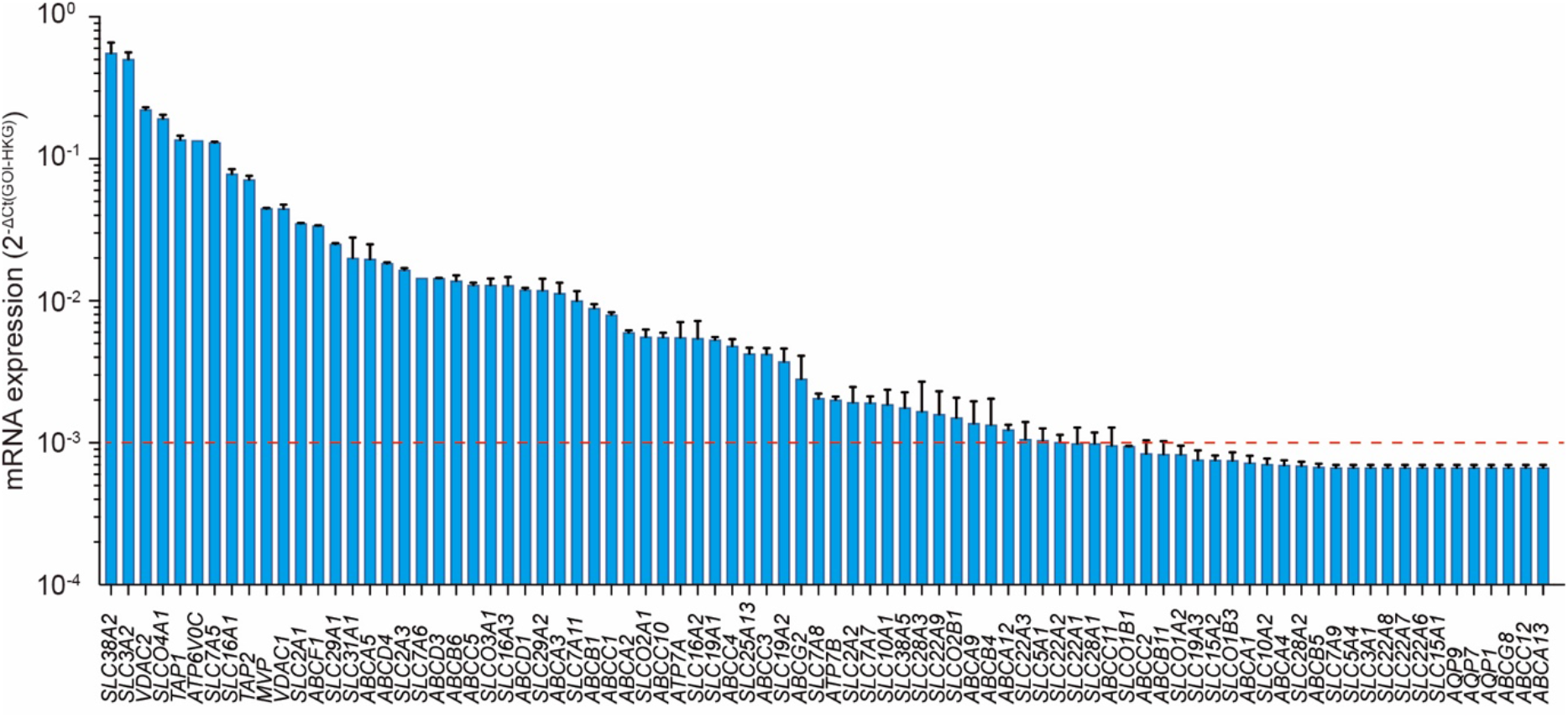
Quantitative analysis of mRNA expression associated with transporters in the Cornea-Chip. The red dashed line indicates the threshold of mRNA expression. Data are presented as means ± S.E.M. (n = 2)

Taken together, these results indicate the ability of our Cornea-Chip model to express major transporters that can mediate the transportation of different nutrients as well as drugs.

### 2.3. Measurement of extracellular metabolites by using Cornea-Chip

To perform metabolomic analysis by using Cornea-Chip, we conducted the procedures shown in **Figure 3A**. In brief, at day 7, the culture medium in the constructed Cornea-Chip was replaced with fresh medium. Next, 1 μL of extracellular culturing medium from both the apical and basolateral sides was collected at 0 h and considered as control. Subsequently, the same amount of sample was collected at 6, 12, 24, and 48 h. Cells were maintained under the same culture conditions to monitor the consumption of nutrients and secretion of metabolites (**Figure 3A** and **Table S3–S5, Supporting Information**). We used calcein AM staining at 48 h to confirm that sample collection did not cause cellular damage (**Figure 3B and C**).

**Figure 3.**
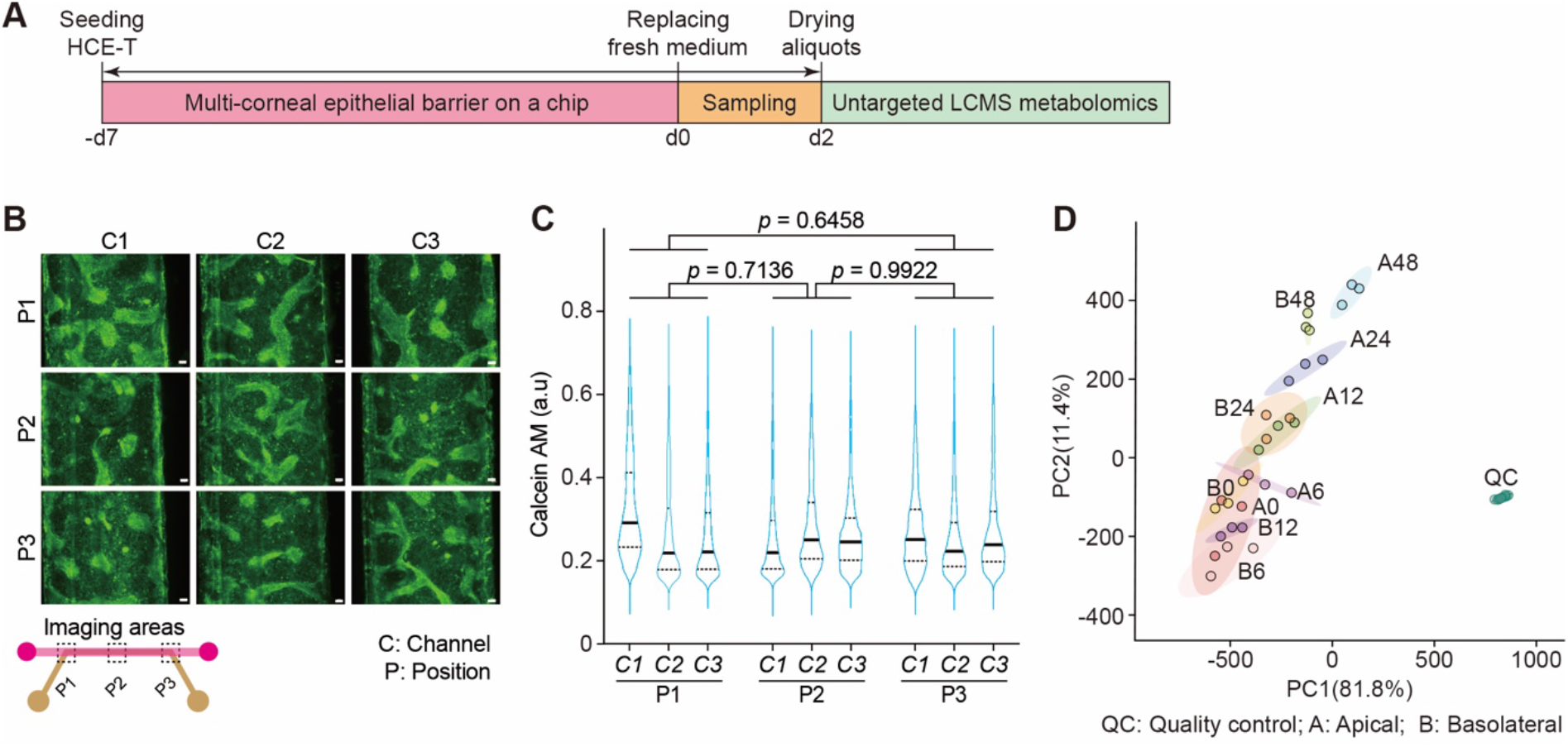
Spatiotemporal analysis of extracellular metabolites from the corneal epithelial barrier on a chip. A) The procedure for collecting and processing extracellular metabolites from the multi-corneal barriers on a chip. B) Cell viability in the microfluidic devices by using calcein-AM staining. HCE-T human corneal epithelial cells in the microfluidic devices after 48 h of metabolite collection. Green, calcein-AM. Scale bar, 100 μm. C) Microscopic signal-cell analysis for evaluating cell viability based on calcein-AM staining shown in B). The analysis was performed on images of 3 independent samples; 4300 cells were randomly selected and analyzed for each sample. Data are represented in the violin plot in which the median of each group is indicated with a scatter line (25^th^ to 75^th^ interquartile range). The *p*-values were determined using Tukey’s multiple comparison test. D) Principal component analysis (PCA) of metabolomics dataset peak areas corrected by channel volume and transformed into cube root values (**Figure S3** - data not corrected by channel volume). The 95% confidence regions are highlighted in different colors. AP: apical, BA: basal, QC: quality controls. Data are derived from 3 biological replicates.

By using our previously developed untargeted LC-MS method for the measurement of extracellular metabolites at the microscale level,^[21]^ we could annotate 104 metabolites at Metabolomics Standard Initiative annotation level 1.^[32]^ Peak areas were used for metabolite semi quantification. All technical internal standards (tISs) showed coefficient of variation (CV) of peak areas of ≤17% across the QC samples and <13% in the study samples (except 5-fluorocytosine, 39.7%). The median CV of all detected metabolites was 9.9% in the QC samples. The median CV of all detected metabolites in the triplicate measurements was 16%, indicating the precision of our untargeted metabolomics workflow in the Cornea-Chip.

Metabolomics data analysis and visualization were conducted by first transforming the peak area value into square roots, with no substantial normalization or scaling (**Figure S2, Supporting Information**). In the principal component analysis (PCA), samples were clustered by all groups indicating systematic metabolite shifts in raw peak area values (**Figure S3, Supporting Information**) as well as in peak area values corrected by the volume of microfluidic channels (**Figure 3D**). QCs and study samples were clearly separated with narrow CIs among the triplicates of each sample in stepwise time-dependent alterations of extracellular metabolites from both apical and basolateral sides. These results indicated that our Cornea-Chip platform facilitated reliable metabolomic analysis and non-invasive spatiotemporal analysis of extracellular metabolites.

### 2.4. Biological pathways and transportation activities

To further explore the corneal metabolic and transporting activities based on the obtained metabolomic profiles, we used the MetaboAnalyst platform.^[33]^ According to the analysis of variance (ANOVA) post-hoc comparison, all annotated metabolites varied significantly (**Figure S4, Supporting Information**). According to the reactome-based biological pathways (*P* < 0.05),^[34]^ 74 out of the 104 metabolites were associated with the transport of nucleosides and free purine and pyrimidine across the plasma membrane, which was mostly associated with SLC transmembrane transporters (SLC28 and SLC29),^[28]^ metabolism-catabolism of nucleotides, purine/pyrimidine catabolism/transportation, metabolism of amino acids and derivatives, and glutathione synthesis/recycling. Moreover, the metabolites showed a significant relationship with a wide range of SLC transporters, including vitamins, organic cation/anion/zwitterion transporters (SLC22),^[35,36]^ as well as ABC transporter family members^[29]^ (**Figure 4A**).

**Figure 4.**
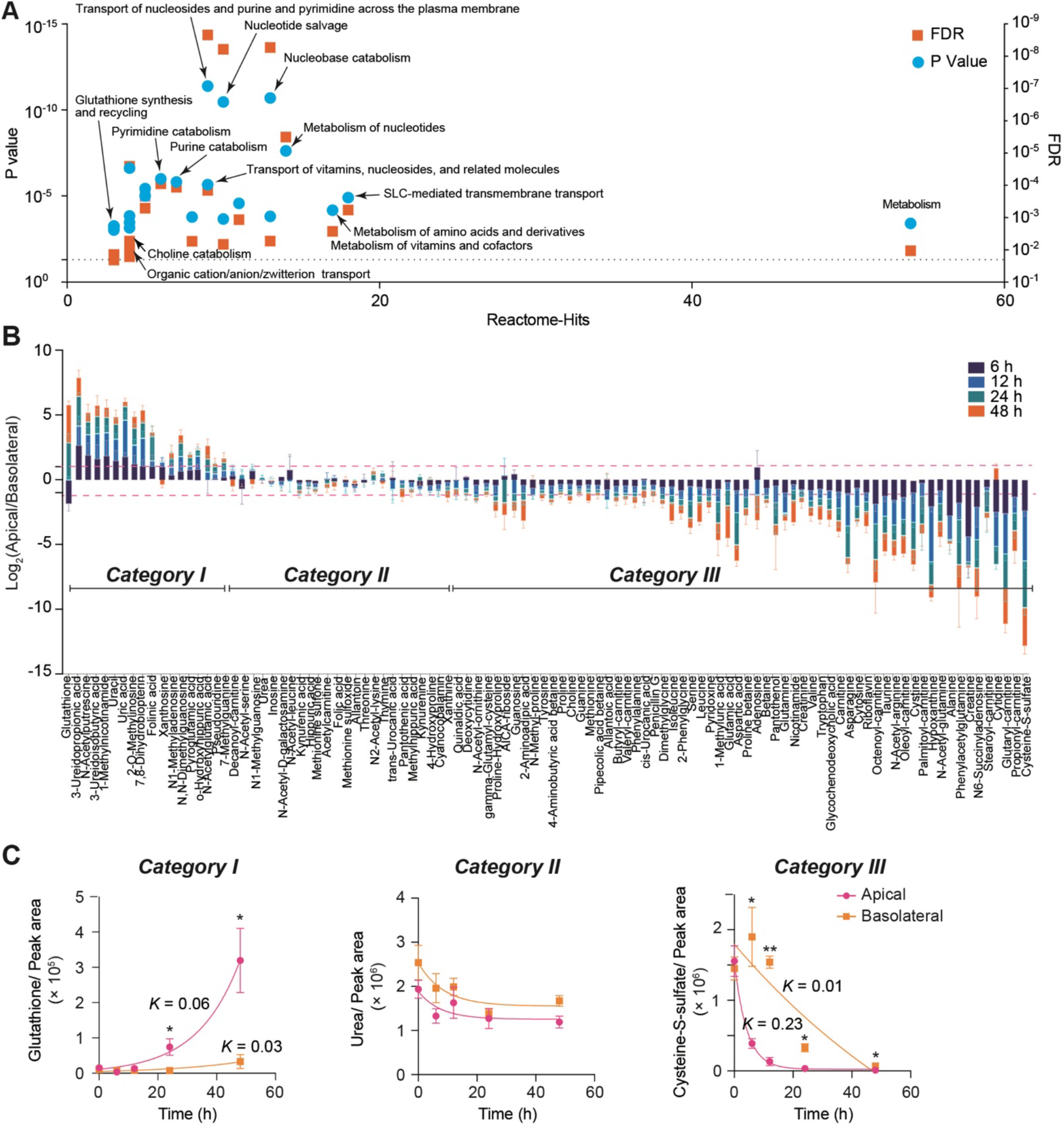
The determination of biological pathways as well as the transportation tendencies of extracellular metabolites. A) Metabolic pathway analysis based on the detection of metabolites found in both the apical and basolateral sides of the corneal epithelial barrier show the prediction hits and statistical significance of *P*-values and false discovery ratio (FDR) values. B) The ratio folds change (log2 apical/basolateral, pink line indicates ±2-fold difference) of metabolites abundances normalized by 0 time point. C) Representative examples of the metabolites in Categories I, II, and III. Peak areas were corrected by the channels volume. Data are presented in triplicates as means ± S.E.M. The secretion and depletion rate (*K*) was obtained using exponential (Malthusian) growth and one phase decay models. \**P*-value <0.05, \*\**P*-value <0.001.

In polarized epithelial barriers, the uptake or secretion of metabolites is known to vary between the apical and basolateral sides. To discriminate these activities, we normalized metabolites by 0 h time point peak areas (considered as control) and then used the log2 ratio of the normalized values of the apical sides divided by those of the basolateral sides as an indicator of the transportation tendency of metabolite secretion (fold change, ≥2; **Figure 4B**).

We observed three different patterns of metabolite transport across the barriers based on the criteria of fold difference of apical/basolateral metabolites: Category I (fold difference, ≥2), in which metabolite secretion was observed in the apical sides or in both the apical and basolateral sides. Category II (−2 < fold difference < 2), where metabolites had no notable secretion or clearance in both the apical and basolateral sides. Category III (fold difference, ≥2), in which nutrient metabolites were gradually depleted from both the apical and basolateral sides with respect to their kinetic rates.

In Category I, glutathione (GSH), uric acid, 3-ureidopropionic acid, 3-ureidoisobutyric acid, 1-methyl nicotinamide, *N*-acetyl putrescine, folinic acid, uracil, 2-*O*-methylinosine, *N*^2^*N*^2^ dimethyl guanosine, *N*^1^-methyl guanosine, and 7-methylguanine showed significant increase (**Figure 4B** and **Figure S5A**). GSH, an endogenous thiol-containing tripeptide, and uric acid derivatives are naturally secreted into human tears and aqueous humor. They are the most abundant antioxidant for their role in cell detoxification of the cornea and conjunctiva.^[37–39]^ The transepithelial transportation of GSH through the apical sides has been reported to be facilitated by ABC transporter family members (ABCC1/C2/C3/C4/C5).^[40–44]^ According to the quantitative RT-PCR results shown in **Figure 2**, the HCE-T cells in the Cornea-Chip expressed ABCC1, ABCC4, and ABCC5, and the former two were expressed on the apical side, and the latter was strongly expressed at the basolateral side.^[30]^ The presence of more efflux transporters in the apical slide of the barrier suggested that the secretion of GSH secretion was greater on the apical side of the barrier (*K* = 0.06 h^−1^) than on the basolateral sides (*K* = 0.03 h^−1^; **Figure 4C**). Moreover, a rapid onset of uric acid secretion was noted on the apical sides at 24 h (**Figure S5A, Supporting Information**). Uric acid and its derivatives are also reported to be effluxed by ABCC4 and ABCG2, which are mainly expressed on the apical sides, as well as SLC22A11/A12 organic anion transporters, which are distributed in both apical and basolateral directions.^[35,45]^ Considering the notable expression of ABCC4 and ABCG2 and basal activities of SLC22A11/A12 in the Cornea-Chip, the secretion kinetics of uric acid might have been regulated by both apical-and basal-related transporters. Taken together, these results indicated that our model could simulate the in vivo secretion of important extracellular metabolites.

As for Category II metabolites, a slight depletion or secretion without any distinction in activity between the apical or basolateral sides. Notably, the interpretation of fold change data does not necessarily suggest that these metabolites are not regulated by cellular transporters such as vitamin and amino acid transporters (**Figure S5B, Supporting Information**). Urea levels slightly decreased, but the depletion rate was not considerably different between the apical and basal sides (**Figure 4C**).

In Category III, metabolites such as cysteine-*S*-sulfate, *N*^6^-succinyladenosine, hypoxanthine, creatine, aspartic acid, phenylacetylglutamine, glutaryl-/octenoyl-/propionyl-/oleoyl-/palmitoyl-carnitine, carnitine, taurine, asparagine, *N*^α^-acetyl arginine, adenosine, and other amino acids were gradually depleted from both the apical and basolateral sides, but with different clearance rates (**Figure 4B** and **Figure S5C, Supporting Information**). Metabolite clearance was quicker in the apical side: the depletion rates (K) of cysteine-*S*-sulfate (**Figure 4C**), hypoxanthine, and oleoyl-carnitine (**Figure S5C, Supporting Information**) were 0.23, 0.25, and 0.2 h^−1^, respectively. In contrast, on the basolateral side, these metabolites showed low K values (0.01, 0.02, and 0.005 h^−1^, respectively; **Figure 4E**). The transportation of amino acids and their derivatives (e.g., cysteine-*S*-sulfate) is known to be mediated by the SLC7 family of transporters,^[46]^ whereas that of nucleoside-related metabolites such as hypoxanthine has been reported to be mediated by nucleoside transporters (e.g., SLC29A2 and SLC29A3) on the apical side of the cornea. Interestingly, RT-PCR results indicated the expression of a wide range of SLC7 family members (SLC7A5/A6/A7/A8/A11) as well as of SLC9A1/A2, which is responsible for the quicker depletion of Category III metabolites, on the apical side of the corneal epithelial barrier.

Remarkably, carnitine and its ester derivatives (glutaryl-/octenoyl-/propionyl-/oleoyl-/palmitoyl-carnitine) were notably depleted on the apical sides of the barrier (**Figure 4B**). The carnitine shuttle is an essential process for delivering fatty acids into the mitochondria for the initiation of ß-oxidation and the tricarboxylic acid cycle, eventually boosting cell energy.^[47]^ The uptake and recycling of carnitine are widely performed by SLC22 transporters (apical direction).^[48,49]^ We showed an essential process of fatty acid transport through the carnitine shuttling system by detecting a wide range of intermediate ester derivatives with different lengths of fatty acids, which are usually difficult to analyze together in vivo owing to their low concentration.^[50]^ The efflux of acylcarnitines from the mitochondria into the cytosol and further into the extracellular space is important during impaired fatty acid oxidation to stop the accumulation of toxic acyl-CoA in the mitochondria; however, the mechanism beyond their release is not yet known.^[51]^

The basolateral activities of metabolite transportation highlighted the potential of our Cornea-Chip in the spatial determination of essential metabolite activities in not only the apical but also the basolateral direction where anatomically layered stromal and corneal endothelial cells exist in the in vivo structure of the human cornea. Such activities have been reported to facilitate the transportation of different nutrients (e.g., glucose) between the corneal epithelium and endothelium.^[52,53]^ Hence, our Cornea-Chip platform can also be used for the elucidation of metabolite molecular signaling among different cells.

## 3. Conclusion

In this study, we introduce multiple corneal barriers on a chip that is feasible for the spatiotemporal collection and analysis of extracellular metabolites. The spatiotemporal determination of extracellular metabolites allowed the investigation of important intracellular biological activities across the corneal epithelium, such as metabolite secretion and depletion. In addition, our approach facilitated the noninvasive prediction of the active transportation sites of extracellular metabolites without requiring additional chemical substrates for a specific transporter. We found that antioxidants such as GSH and uric acid could be secreted from the corneal cells in the Cornea-Chip simulating similar vivo secretion conditions. These metabolites are attractive biomarkers for measuring the oxidative stress level in cells and can thus be considered as indicators of cell toxicity during the obstruction of corneal homeostasis. These findings can form the basis for the further investigation of the secretion and transportation of xenogenous and endogenous compounds across the corneal epithelium in the future.

## 4. Experimental Section

### Microfluidic device fabrication

The microfluidic device was fabricated using stereolithographic 3D-printing techniques and solution cast-molding processes.^[15,54]^ In brief, the mold for microfluidic channels was produced using a 3D printer (Keyence Corporation, Osaka, Japan). Two molds were fabricated: the upper and lower blocks. Each block contained four chambers (15 mm length, 1.5 mm width, and 0.5 mm height). Before use, the surfaces of the molds were coated with trichloro(1H,1H,2H,2H-perfluorooctyl) silane (Sigma-Aldrich, St. Louis, MO, USA). Sylgard 184 PDMS two-part elastomer (ratio of pre-polymer to curing agent, 10:1; Dow Corning Corporation, Midland, MI, USA) was mixed, poured into the molds to produce 4-mm and 0.5-mm thick PDMS upper and lower layers, respectively, and degassed using a vacuum desiccator for 1 h. The PDMS of the lower block was fixed on a glass slide and then cured in an oven at 80°C for 24 h. After curing, the PDMS was removed from the molds, trimmed, and cleaned. A clear PET membrane was fixed on each chamber of the lower PDMS block. Both the PDMS blocks were treated with corona plasma (Shinko Denki, Inc., Oosaka, Japan) and bonded together by baking in an oven at 80°C. Scanning electron micrographs were obtained using a JCM-5000 microscope (JEOL Ltd., Tokyo, Japan) at 10 kV. Before imaging, a 5-nm thick platinum layer was sputtered on the samples (MSP 30 T; Shinku Device, Sagamihara, Japan).

### Human corneal epithelial cell culture

HCE-T cells were provided by RIKEN Bioresource Research Centre (Ibraki, Japan). Cells were cultured in DMEM/F12 supplemented with 5% (v/v) fetal bovine serum, 5 μg mL^−1^ insulin, 10 ng mL^−1^ human epithelial growth factor, and 0.5% dimethyl sulfoxide. The cells were passaged with trypsin-EDTA (0.25–0.02%) solution at a 1:4 subculture ratio.

### Human corneal epithelial barrier construction in the microfluidic device

Before use, the microfluidic cell culture devices were placed under ultraviolet light in a biosafety cabinet for 30 min. The microfluidic channels were washed with DMEM/F12. Cells were harvested using trypsin and collected in a 15 mL tube. Following centrifugation, the cells were resuspended in DMEM/F12 medium and introduced into the upper channel of the microfluidic devices via a cell inlet with a cross-sectional area of 0.23 cm^2^ at a density of 1 × 10^6^ cells mL^−1^. The lower receiver channel was filled with DMEM/F12 only. The microfluidic devices were then placed in a humidified incubator at 37°C with 5% CO2 for 7 days. The medium in each chamber was periodically changed every 24 h.

### Cell viability assay

Cell viability was assessed by live staining with calcein AM (Dojindo Molecular Technologies, Inc.). In brief, cells were incubated with calcein AM at a final concentration of 10 μg mL^−1^ in DMEM/F12 medium at 37°C for 60 min. The cells were then washed twice with PBS and subjected to microscopy imaging.

### Immunofluorescence and microscopy imaging

For immunostaining, cells were fixed with 4% paraformaldehyde in PBS for 25 min at 25°C and then permeabilized with 0.5% Triton X-100 in PBS for 10 min at 25°C. Subsequently, the cells were blocked with blocking buffer [5% (v/v) normal goat serum, 5% (v/v) normal donkey serum, 3% (w/v) bovine serum albumin, 0.1% (v/v) Tween-20] at 4°C for 24 h and then incubated at 4°C overnight with the primary antibodies in blocking buffer (**Table S1, Supporting Information**). Cells were then incubated at 37°C for 60 min with a secondary antibody (Alexa Fluor 594 donkey anti-rabbit IgG and Alexa Fluor 594 donkey anti-mouse IgG, 1:1000; Jackson ImmunoResearch, West Grove, PA, USA) in blocking buffer before final incubation with 4,6-diamidino-2-phenylindole (DAPI) or anti-phalloidin for F-actin at 25°C. For imaging, we used a Nikon ECLIPSE Ti inverted fluorescence microscope equipped with a CFI plan fluor (10×/0.30 N.A. objective lens; Nikon, Tokyo, Japan). Z-scans were obtained using an Andor Multi-model Fast Confocal Microscope System Dragonfly (Oxford Instruments, Abingdon, UK). Images were then analyzed using ImageJ software (National Institute of Health, Maryland, USA) or CellProfiler software (Version 3.1.8; Broad Institute of Harvard and MIT, USA^[55]^). *Quantitative RT-PCR Array*: Total RNA from microfluidic channels was purified using an RNeasy Mini Kit (Qiagen). Next, 0.3 μg of total RNA was reverse-transcribed using the RT First-strand Kit (Qiagen). The cDNA obtained in solution was mixed with Power SYBR Green PCR MasterMix (Life Technologies) and introduced into the Drug Transporter-specific RT2 Array (Qiagen, Cat. no. 330231 PAHS-070ZA) in a 96-well format, according to the manufacturer’s instructions. PCR conditions were as follows: initial incubation at 95°C for 10 min, followed by 40 cycles of 95°C for 15 s and then 60°C for 3 min; an Applied Biosystems 7300 Real-Time PCR system (Life Technologies) was used for the PCRs. The mRNA repression was performed using the 2^−ΔCT^ method, where ΔCT = CT (gene of interest; GOI) - CT (housekeeping genes; HKG).

### Extracellular culture medium collection for untargeted metabolomics

After the HCE-T cells were cultured for 7 days, the culture medium was replaced with fresh medium in both the apical and basolateral channels of the microfluidic device. Microfluidic devices were then placed at 37°C with 5% CO2. At 0, 6, 12, 24 and 48 h time points, a sample consisting of 1 μL of extracellular culturing medium from both the apical and basolateral channels was collected. The collected samples were dried in a vacuum incubator for 3 h at 25°C and then preserved at −80°C.

### Untargeted LC-MS metabolomics

Tubes containing 1 μL of dried sample were thawed and 100 μL of water:acetonitrile (1:9, v/v) mixture containing 5 tISs was added (**Table S2, Supporting Information**). For the preparation of QC samples, 0 h cell culture medium sample was used from our previous study.^[21]^ For each QC sample, 1 μL of cell culture medium was evaporated and processed together. After resuspension, the samples were centrifuged at 4°C for 15 min at 20000 *g*. Next, 40 μL of the supernatant was transferred to a 96-well 0.2 mL PCR plate (PCR-96-MJ; BMBio, Tokyo, Japan). The plate was sealed with a pierceable seal (4titude; Wotton, UK) for 3 s at 180°C by using a plate sealer (BioRad PX-1; CA, USA) and maintained at 4°C during the LC-MS measurement. The injection volume was 10 μL. The LC-MS method has been described previously.^[56–58]^ In brief, metabolite separation was achieved on an Agilent 1290 Infinity II system by using SeQuant ZIC-HILIC (Merck, Darmstadt, Germany) column by using a 12 min gradient of acidified acetonitrile and water. Data were acquired on an Agilent 6550 Q-TOF-MS system with a mass range of 40-1200 m/z in the positive all ion fragmentation mode, including 3 sequential experiments at alternating collision energies: one full scan at 0 eV, followed by one MS/MS scan at 10 eV, and then one MS/MS scan at 30 eV. The data acquisition rate was 6 scans/s. Data were converted to mzML format by using Proteowizard and processed using MS-DIAL version 4.38^[59,60]^ (detailed parameters are shown in **Tables S4 and S5, Supporting Information**). An in-house MS2 spectral library containing experimental MS2 spectra and retention times (RTs) for 391 compounds obtained from standards^[56,58,61]^ was used to annotate the detected compounds on the basis of 3 criteria: (i) accurate mass (AM) match (tolerance: 0.01 Da), (ii) RT match (tolerance: 1 min), and (iii) MS2 spectrum match (similarity, >80%). The MS2 similarity was scored as a simple dot product without any weighting (at least two MS2 peaks matched with the reference spectra). The MS2 similarities with reference spectra were matched to any of the CorrDec^[62]^ or MS2Dec^[60]^ deconvoluted MS2 spectra of the three collision energies (0, 10, and 30 eV). Peak areas exported from MS-DIAL were used for metabolite semiquantification. Metabolites with CV of <30% in QC samples or D-ratio of <50% were used for further analyses.^[63]^ The dataset has been deposited to the EMBL-EBI MetaboLights repository with the identifier MTBLS2274.^[64]^

## Supporting Information

Supporting Information is available from the Wiley Online Library or from the author.

## Acknowledgements

R.A. and R.C. contributed equally to the project. Funding was generously provided by the Japan Society for the Promotion of Science (JSPS; 20K20168, 17H02083, 18KK0306, and 19H02572), Japan Agency for Medical Research and Development (AMED; 17937667), Kyoto University GAP fund program (207010), and LiaoNing Revitalization Talents Program (XLYC1902061). WPI-iCeMS is supported by the World Premier International Research Centre Initiative (WPI), MEXT, Japan. We acknowledge the support from the Gunma University Initiative for Advanced Research (GIAR). We also acknowledge the iCeMS analysis center for providing the imaging facility.

## Supporting Information

**Table S1.** List of reagents and resources for fluorescent immunocytochemistry

| **Item** | **Maker** | **Catalog**<br>**number** | **Dilution** |
| --- | --- | --- | --- |
| ZO-1 antibody,<br>Rabbit | Thermo Fisher | 61-7300 | IF 1:50 |
| P-gp antibody,<br>Rabbit | Abcam | ab129450 | IF 1:50 |
| CK12 antibody,<br>Mouse | Santa Cruz<br>Biotechnology | sc-515882 | IF 1:200 |
| Phalloidin-iFluor<br>594 Reagent | Abcam | ab176757 |  |
| Prism Software | Version 8; La Jolla<br>California, USA |  |  |
| CellProfiler<br>Software | Version 3.1.8; Broad<br>Institute of Harvard and<br>MIT, USA |  |  |

**Table S2.** List of transporter genes for quantitative RT-PCR

| **RefSeq ID** | **Symbol** | **Description** |
| --- | --- | --- |
| NM_005502 | ABCA1 | ATP-binding cassette, sub-family A (ABC1), member 1 |
| NM_173076 | ABCA12 | ATP-binding cassette, sub-family A (ABC1), member 12 |
| NM_152701 | ABCA13 | ATP-binding cassette, sub-family A (ABC1), member 13 |
| NM_001606 | ABCA2 | ATP-binding cassette, sub-family A (ABC1), member 2 |
| NM_001089 | ABCA3 | ATP-binding cassette, sub-family A (ABC1), member 3 |
| NM_000350 | ABCA4 | ATP-binding cassette, sub-family A (ABC1), member 4 |
| NM_018672 | ABCA5 | ATP-binding cassette, sub-family A (ABC1), member 5 |
| NM_080283 | ABCA9 | ATP-binding cassette, sub-family A (ABC1), member 9 |
| NM_000927 | ABCB1 | ATP-binding cassette, sub-family B (MDR/TAP), member 1 |
| NM_003742 | ABCB11 | ATP-binding cassette, sub-family B (MDR/TAP), member 11 |
| NM_000443 | ABCB4 | ATP-binding cassette, sub-family B (MDR/TAP), member 4 |
| NM_178559 | ABCB5 | ATP-binding cassette, sub-family B (MDR/TAP), member 5 |
| NM_005689 | ABCB6 | ATP-binding cassette, sub-family B (MDR/TAP), member 6 |
| NM_004996 | ABCC1 | ATP-binding cassette, sub-family C (CFTR/MRP), member 1 |
| NM_033450 | ABCC10 | ATP-binding cassette, sub-family C (CFTR/MRP), member 10 |
| NM_032583 | ABCC11 | ATP-binding cassette, sub-family C (CFTR/MRP), member 11 |
| NM_033226 | ABCC12 | ATP-binding cassette, sub-family C (CFTR/MRP), member 12 |
| NM_000392 | ABCC2 | ATP-binding cassette, sub-family C (CFTR/MRP), member 2 |
| NM_003786 | ABCC3 | ATP-binding cassette, sub-family C (CFTR/MRP), member 3 |
| NM_005845 | ABCC4 | ATP-binding cassette, sub-family C (CFTR/MRP), member 4 |
| NM_005688 | ABCC5 | ATP-binding cassette, sub-family C (CFTR/MRP), member 5 |
| NM_000033 | ABCD1 | ATP-binding cassette, sub-family D (ALD), member 1 |
| NM_002858 | ABCD3 | ATP-binding cassette, sub-family D (ALD), member 3 |
| NM_005050 | ABCD4 | ATP-binding cassette, sub-family D (ALD), member 4 |
| NM_001090 | ABCF1 | ATP-binding cassette, sub-family F (GCN20), member 1 |
| NM_004827 | ABCG2 | ATP-binding cassette, sub-family G (WHITE), member 2 |
| NM_022437 | ABCG8 | ATP-binding cassette, sub-family G (WHITE), member 8 |
| NM_198098 | AQP1 | Aquaporin 1 (Colton blood group) |
| NM_001170 | AQP7 | Aquaporin 7 |
| NM_020980 | AQP9 | Aquaporin 9 |
| NM_001694 | ATP6V0C | ATPase, H<sup>+</sup> transporting, lysosomal 16 kDa, V0 subunit c |
| NM_000052 | ATP7A | ATPase, Cu<sup>++</sup> transporting, alpha polypeptide |
| NM_000053 | ATP7B | ATPase, Cu<sup>++</sup> transporting, beta polypeptide |
| NM_017458 | MVP | Major vault protein |
| NM_003049 | SLC10A1 | Solute carrier family 10 (sodium/bile acid cotransporter family), member 1 |
| NM_000452 | SLC10A2 | Solute carrier family 10 (sodium/bile acid cotransporter family), member 2 |
| NM_005073 | SLC15A1 | Solute carrier family 15 (oligopeptide transporter), member 1 |
| NM_021082 | SLC15A2 | Solute carrier family 15 (H<sup>+</sup>/peptide transporter), member 2 |
| NM_003051 | SLC16A1 | Solute carrier family 16, member 1 (monocarboxylic acid transporter 1) |
| NM_006517 | SLC16A2 | Solute carrier family 16, member 2 (monocarboxylic acid transporter 8) |
| NM_004207 | SLC16A3 | Solute carrier family 16, member 3 (monocarboxylic acid transporter 4) |
| NM_194255 | SLC19A1 | Solute carrier family 19 (folate transporter), member 1 |
| NM_006996 | SLC19A2 | Solute carrier family 19 (thiamine transporter), member 2 |
| NM_025243 | SLC19A3 | Solute carrier family 19, member 3 |
| NM_003057 | SLC22A1 | Solute carrier family 22 (organic cation transporter), member 1 |
| NM_003058 | SLC22A2 | Solute carrier family 22 (organic cation transporter), member 2 |
| NM_021977 | SLC22A3 | Solute carrier family 22 (extraneuronal monoamine transporter), member 3 |
| NM_004790 | SLC22A6 | Solute carrier family 22 (organic anion transporter), member 6 |
| NM_006672 | SLC22A7 | Solute carrier family 22 (organic anion transporter), member 7 |
| NM_004254 | SLC22A8 | Solute carrier family 22 (organic anion transporter), member 8 |
| NM_080866 | SLC22A9 | Solute carrier family 22 (organic anion transporter), member 9 |
| NM_014251 | SLC25A13 | Solute carrier family 25, member 13 (citrin) |
| NM_004213 | SLC28A1 | Solute carrier family 28 (sodium-coupled nucleoside transporter), member 1 |
| NM_004212 | SLC28A2 | Solute carrier family 28 (sodium-coupled nucleoside transporter), member 2 |
| NM_022127 | SLC28A3 | Solute carrier family 28 (sodium-coupled nucleoside transporter), member 3 |
| NM_004955 | SLC29A1 | Solute carrier family 29 (nucleoside transporters), member 1 |
| NM_001532 | SLC29A2 | Solute carrier family 29 (nucleoside transporters), member 2 |
| NM_006516 | SLC2A1 | Solute carrier family 2 (facilitated glucose transporter), member 1 |
| NM_000340 | SLC2A2 | Solute carrier family 2 (facilitated glucose transporter), member 2 |
| NM_006931 | SLC2A3 | Solute carrier family 2 (facilitated glucose transporter), member 3 |
| NM_001859 | SLC31A1 | Solute carrier family 31 (copper transporters), member 1 |
| NM_018976 | SLC38A2 | Solute carrier family 38, member 2 |
| NM_033518 | SLC38A5 | Solute carrier family 38, member 5 |
| NM_000341 | SLC3A1 | Solute carrier family 3 (cystine, dibasic and neutral amino acid transporters, activator of cystine, dibasic and neutral amino acid transport), member 1 |
| NM_002394 | SLC3A2 | Solute carrier family 3 (activators of dibasic and neutral amino acid transport), member 2 |
| NM_000343 | SLC5A1 | Solute carrier family 5 (sodium/glucose cotransporter), member 1 |
| NM_014227 | SLC5A4 | Solute carrier family 5 (low affinity glucose cotransporter), member 4 |
| NM_014331 | SLC7A11 | Solute carrier family 7 (anionic amino acid transporter light chain, xc-system), member 11 |
| NM_003486 | SLC7A5 | Solute carrier family 7 (amino acid transporter light chain, L system), member 5 |
| NM_003983 | SLC7A6 | Solute carrier family 7 (amino acid transporter light chain, y + L system), member 6 |
| NM_003982 | SLC7A7 | Solute carrier family 7 (amino acid transporter light chain, y + L system), member 7 |
| NM_182728 | SLC7A8 | Solute carrier family 7 (amino acid transporter light chain, L system), member 8 |
| NM_014270 | SLC7A9 | Solute carrier family 7 (glycoprotein-associated amino acid transporter light chain, bo,+ system), member 9 |
| NM_021094 | SLCO1A2 | Solute carrier organic anion transporter family, member 1A2 |
| NM_006446 | SLCO1B1 | Solute carrier organic anion transporter family, member 1B1 |
| NM_019844 | SLCO1B3 | Solute carrier organic anion transporter family, member 1B3 |
| NM_005630 | SLCO2A1 | Solute carrier organic anion transporter family, member 2A1 |
| NM_007256 | SLCO2B1 | Solute carrier organic anion transporter family, member 2B1 |
| NM_013272 | SLCO3A1 | Solute carrier organic anion transporter family, member 3A1 |
| NM_016354 | SLCO4A1 | Solute carrier organic anion transporter family, member 4A1 |
| NM_000593 | TAP1 | Transporter 1, ATP-binding cassette, sub-family B (MDR/TAP) |
| NM_000544 | TAP2 | Transporter 2, ATP-binding cassette, sub-family B (MDR/TAP) |
| NM_003374 | VDAC1 | Voltage-dependent anion channel 1 |
| NM_003375 | VDAC2 | Voltage-dependent anion channel 2 |
| NM_001101 | ACTB | Actin, beta (HKG) |
| NM_004048 | B2M | Beta-2-microglobulin (HKG) |
| NM_002046 | GAPDH | Glyceraldehyde-3-phosphate dehydrogenase (HKG) |
| NM_000194 | HPRT1 | Hypoxanthine phosphoribosyltransferase 1(HKG) |
| NM_001002 | RPLP0 | Ribosomal protein, large, P0(HKG) |

**Table S3.**
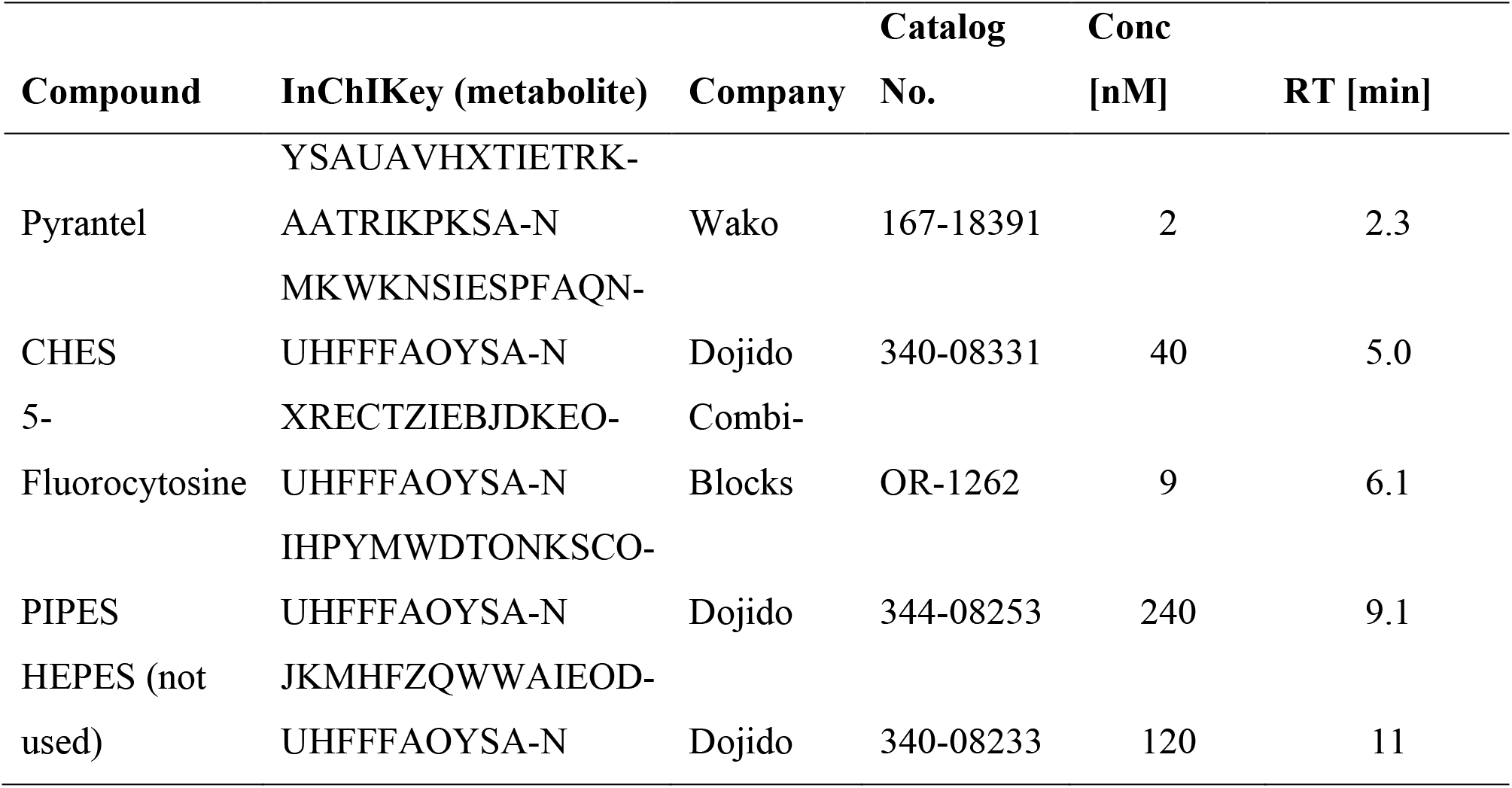
List of internal standards

**Table S4.** MSDIAL experiment file

| **ID** | **MS Type** | **Start m/z** | **End m/z** | **Name** | **Collision energy** | **Deconvolution target (0: No, 1: Yes)** |
| --- | --- | --- | --- | --- | --- | --- |
| 0 | ALL | 40 | 1200 | 10 eV | 10 | 1 |
| 1 | ALL | 40 | 1200 | 30 eV | 30 | 1 |
| 2 | SCAN | 40 | 1200 | 0 eV | 0 | 1 |

**Table S5.**
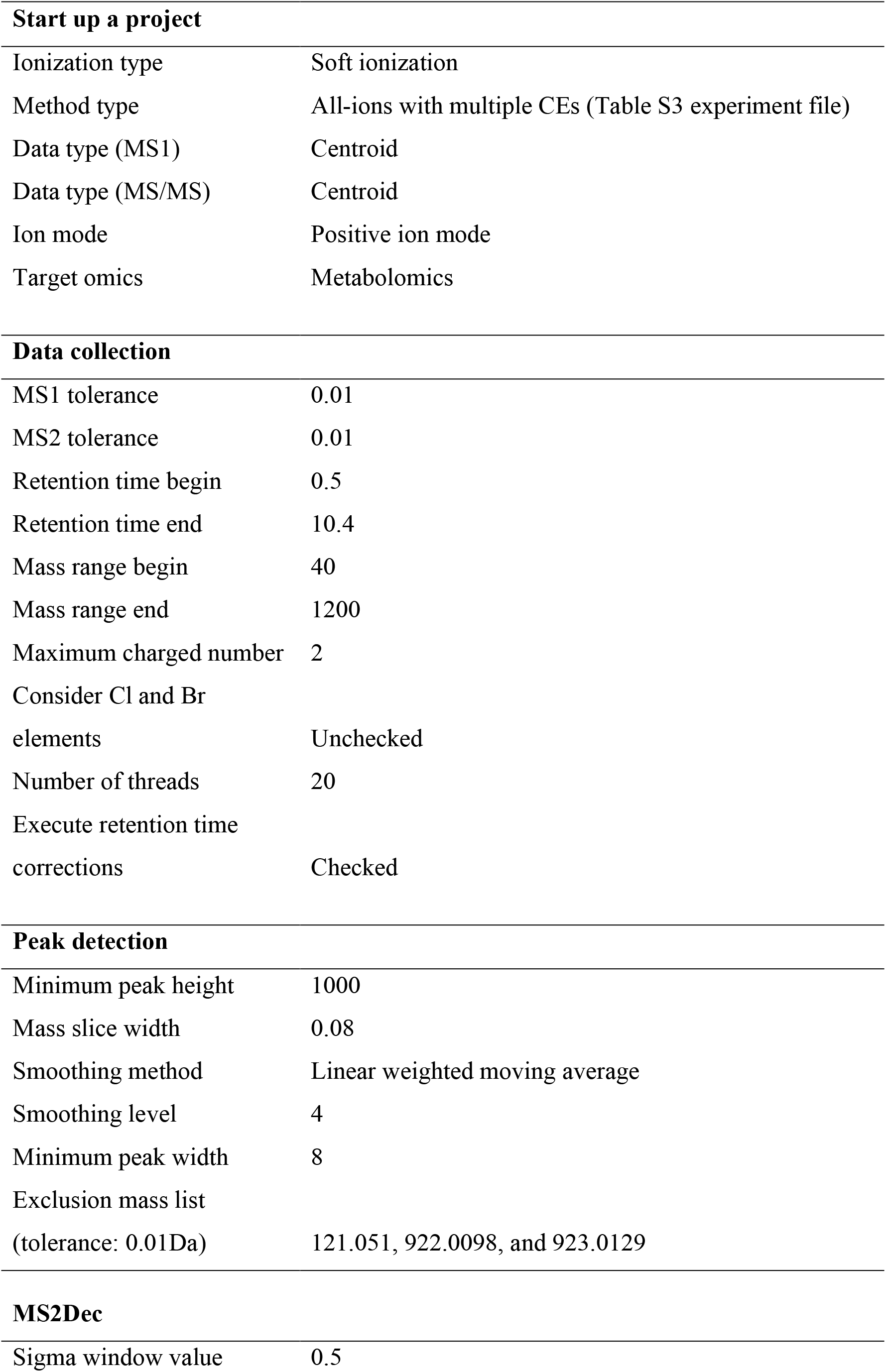
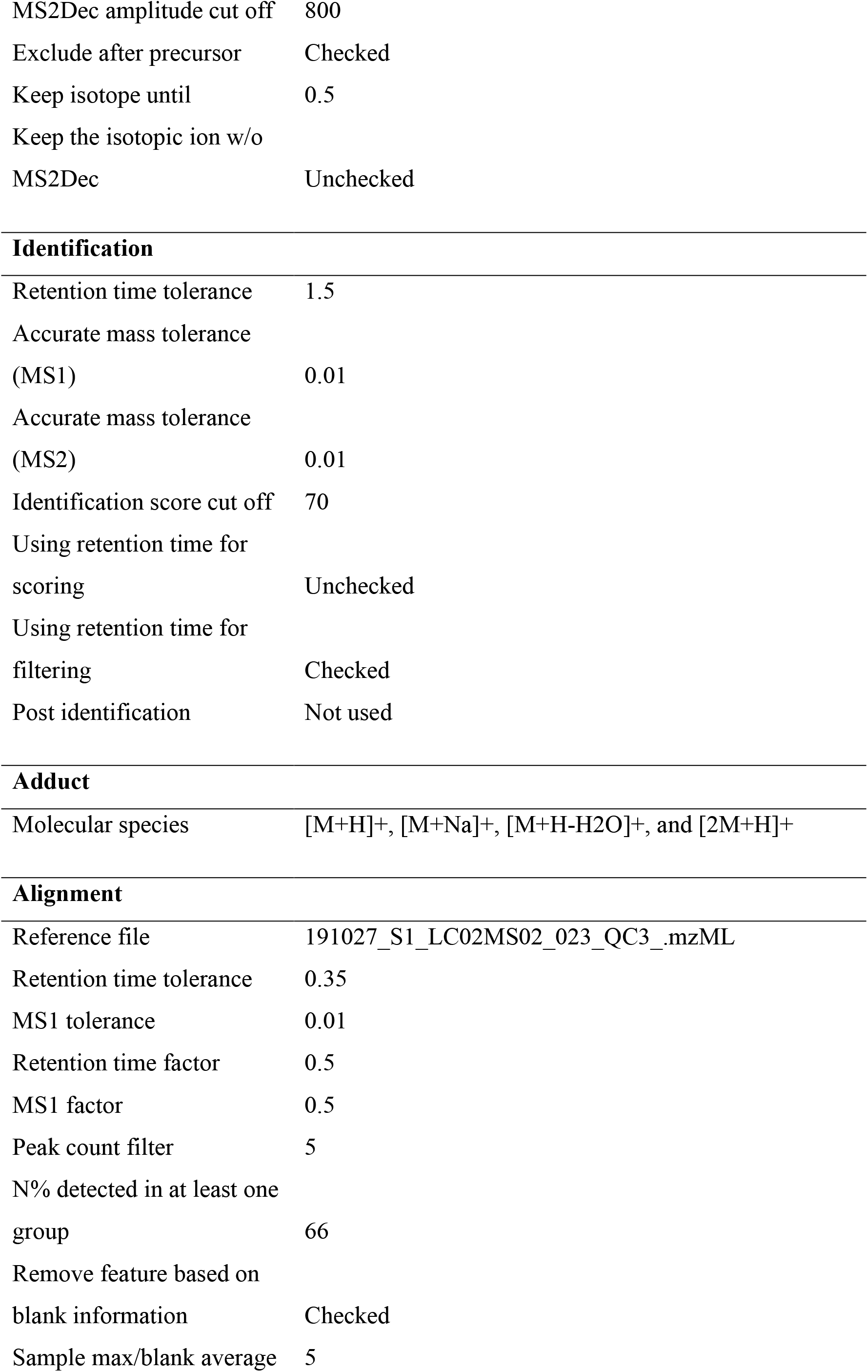
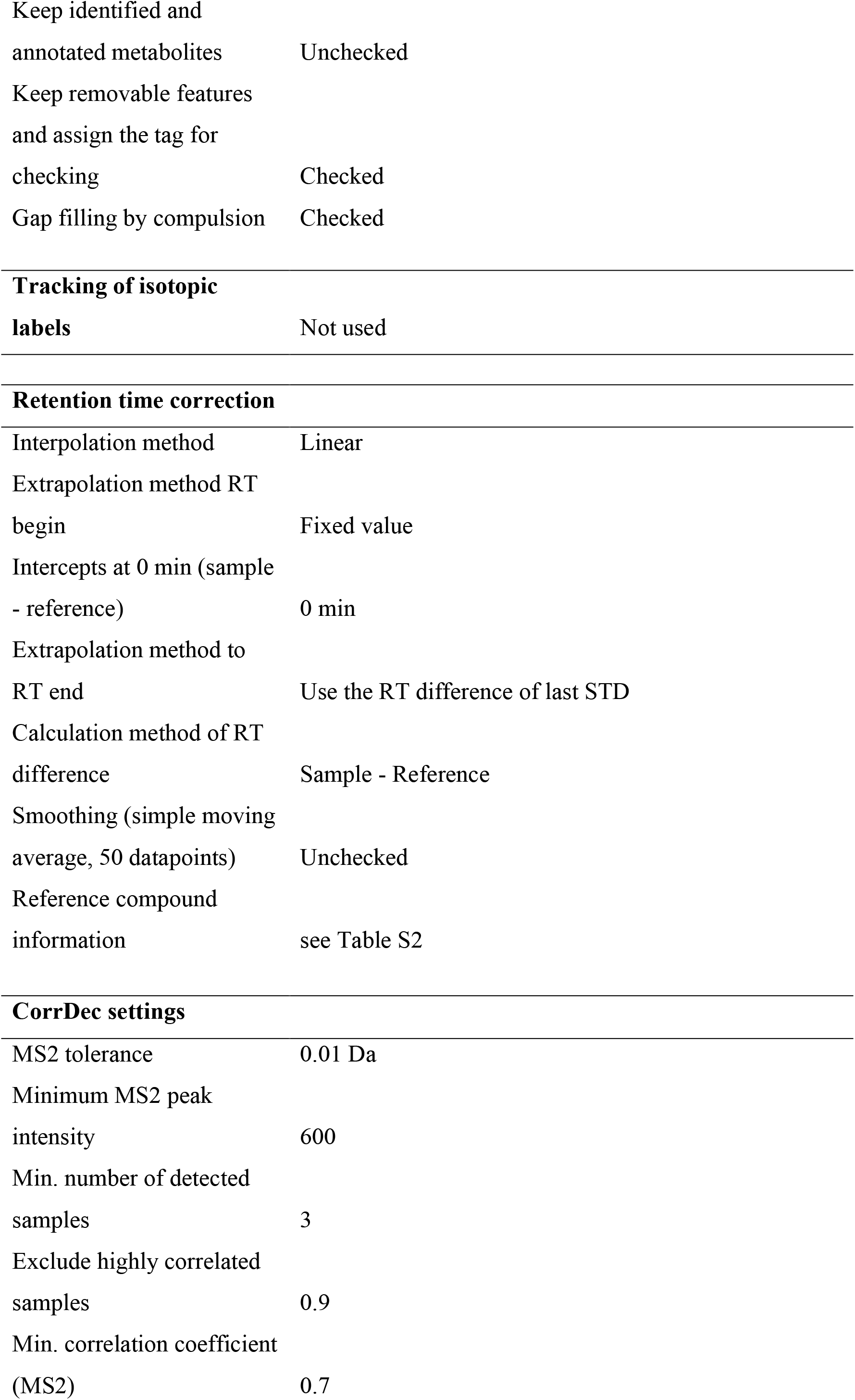
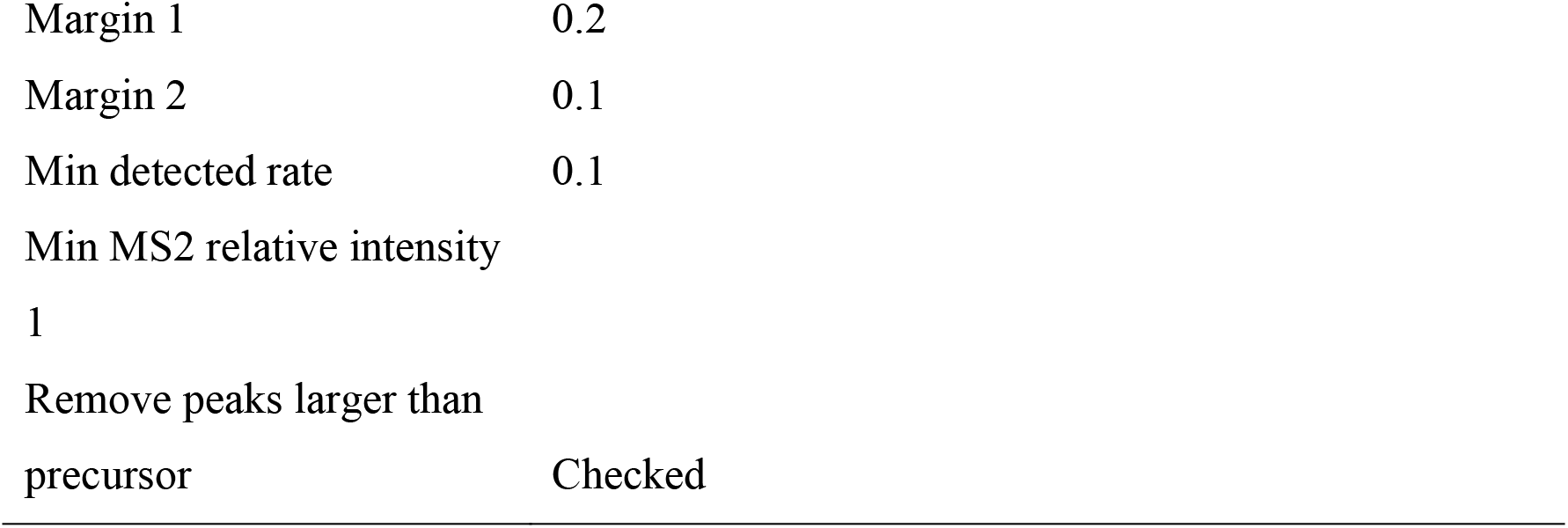
MSDIAL project setting

**Figure S1.**
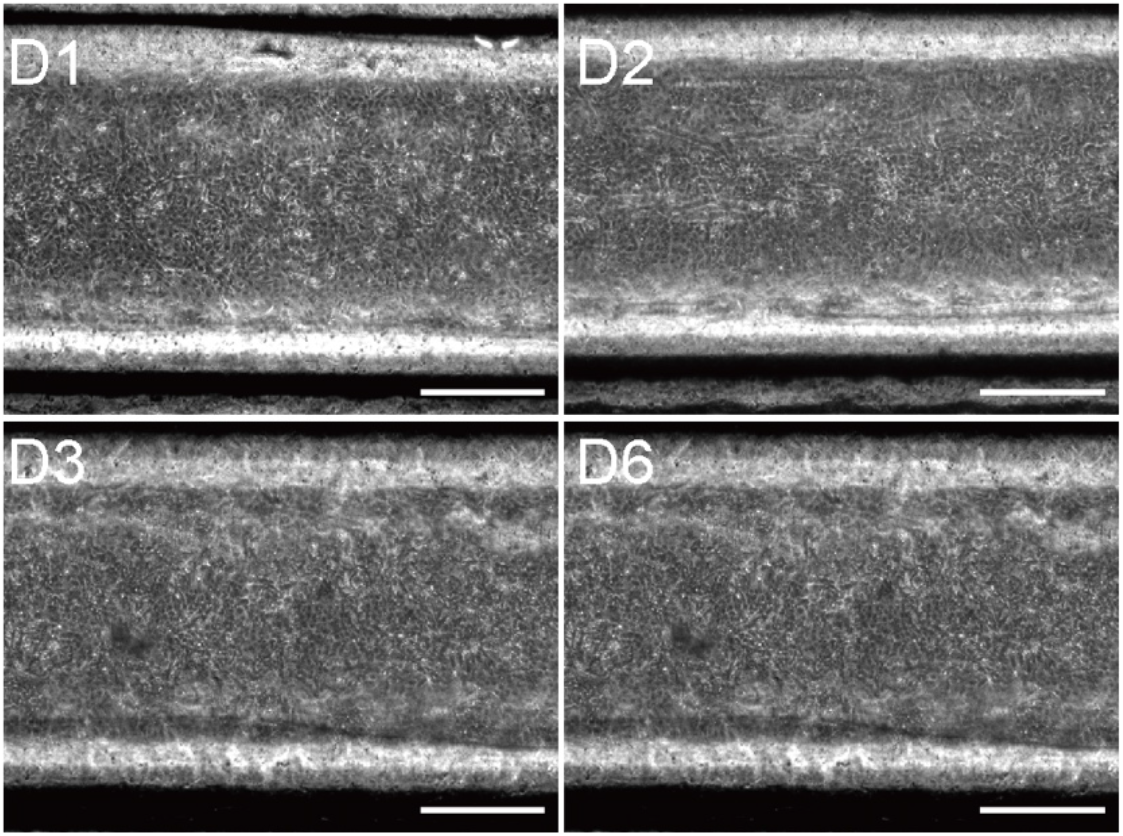
Microscopic bright field images of HCE-T cells in the Cornea-Chip. Scale bar, 500 μm.

**Figure S2.**
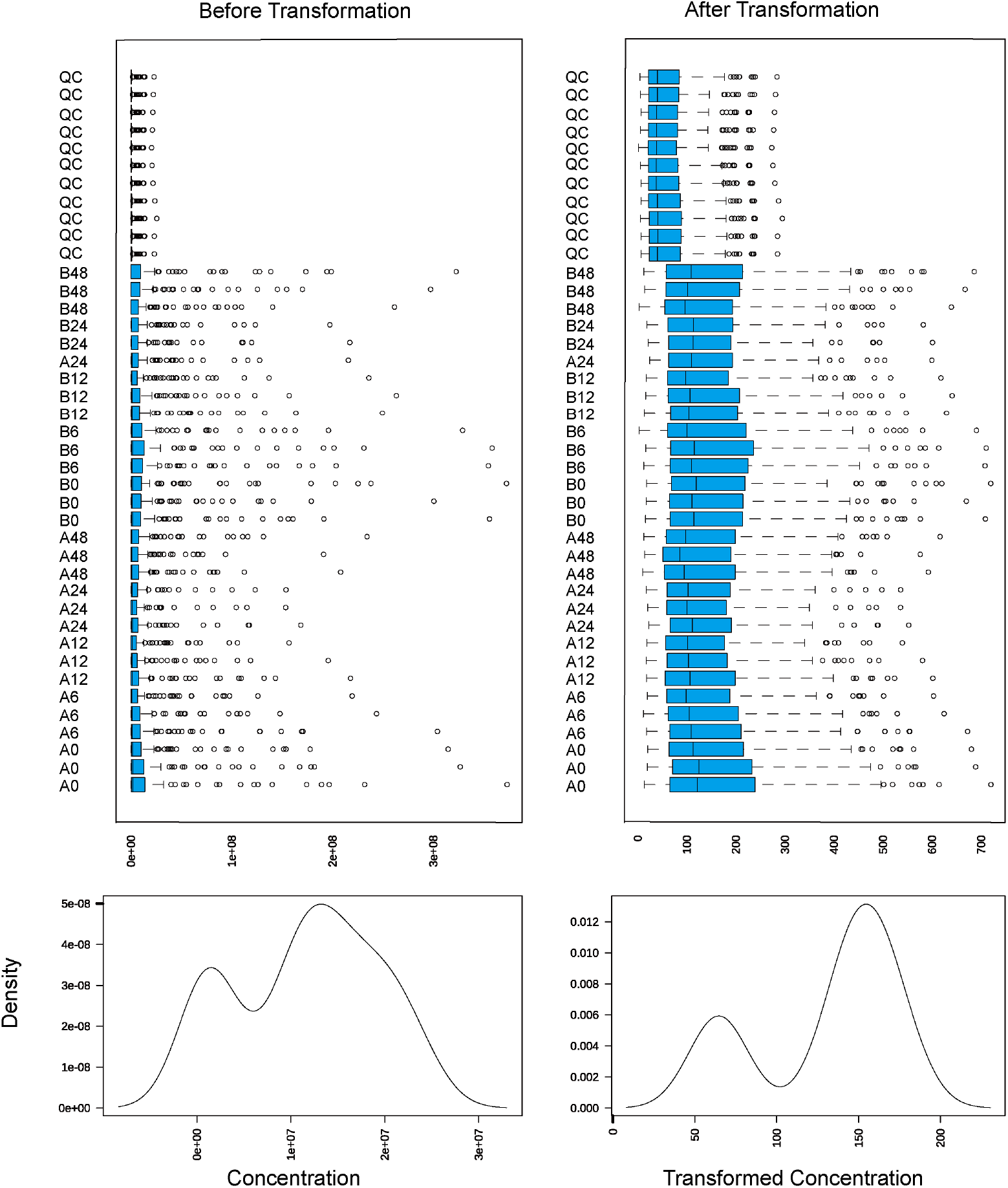
Data processing and transformation. Data were transformed by considering the square root of sample values. Boxplots with the median of each group (25^th^–75^th^ interquartile range, the whiskers extend between maximum and minimum values). Q: quality control samples. A: Apical. B: Basolateral.

**Figure S3.**
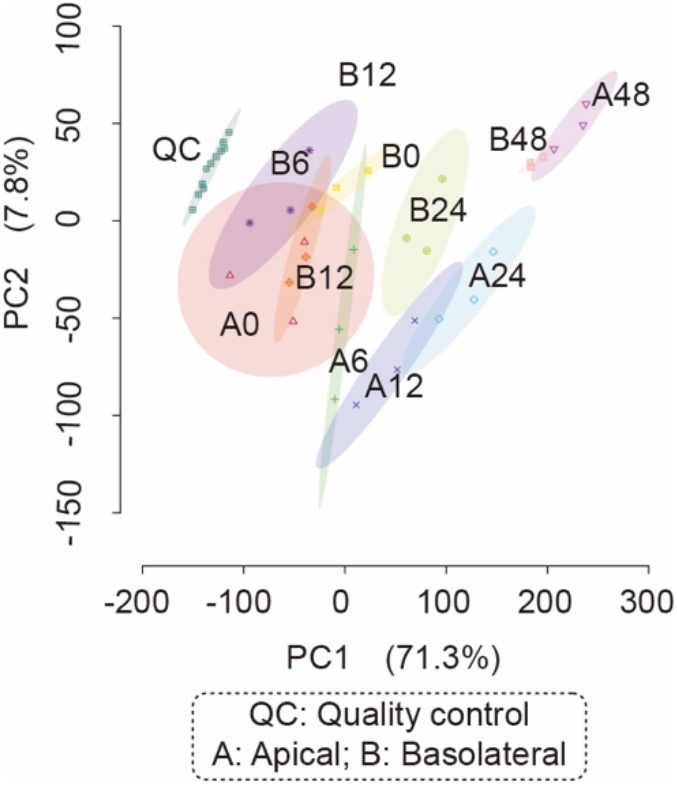
Principal component analysis (PCA) of original value samples and quality controls. Data were transformed into cube root values. The 95% confidence regions are highlighted in different colors. AP: apical, BA: basolateral, QC: quality controls.

**Figure S4.**
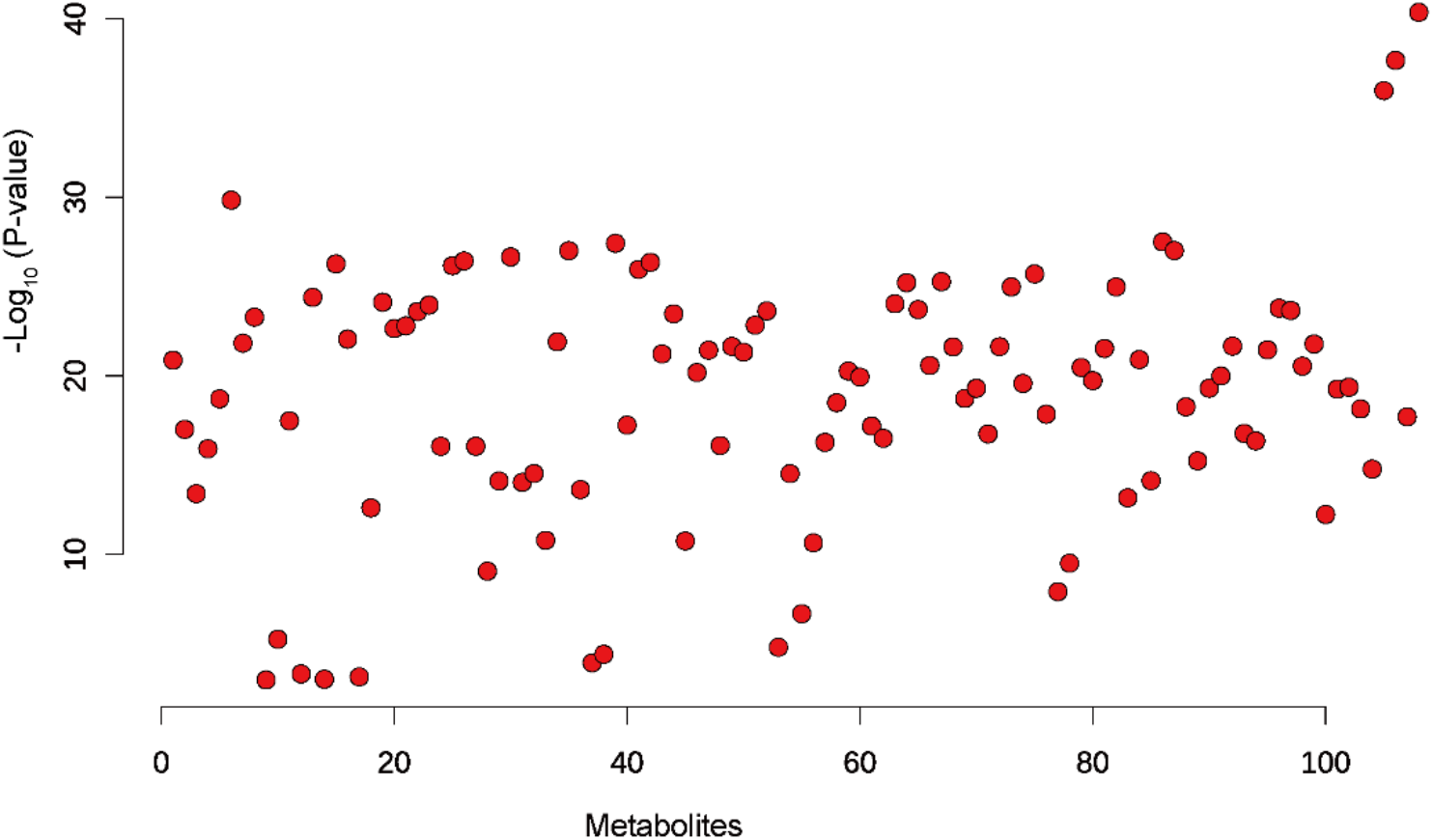
One-way ANOVA and post-hoc test analysis of all samples and QC. *P*-value < 0.05

**Figure S5.**
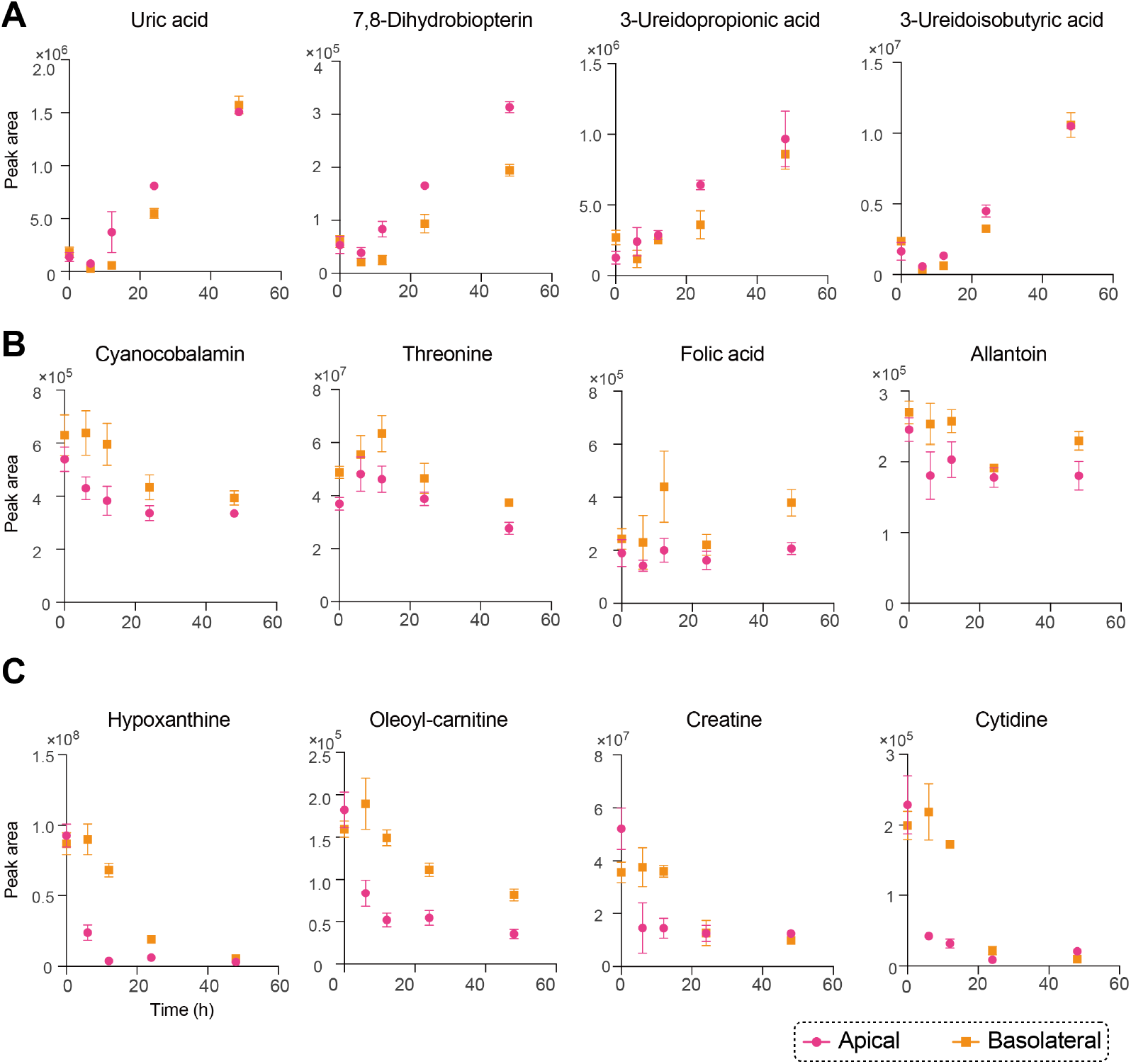
Selected metabolite activities across the Cornea-chip. A–C) Metabolites associated with categories I, II, and III, respectively. Peak areas were corrected by the channels volume. Data are presented in triplicates as means ± S.E.M.

## References

[1] J. Nirmal, S. B. Singh, N. R. Biswas, V. Thavaraj, R. V. Azad, T. Velpandian, Eye 2013, 27, 1196.

[2] C. D. Xiang, M. Batugo, D. C. Gale, T. Zhang, J. Ye, C. Li, S. Zhou, E. Y. Wu, E. Y. Zhang, Drug Metab. Dispos. 2009, 37, 992.

[3] J. Čejková, M. Vejražka, J. Pláteník, S. Štípek, Exp. Gerontol. 2004, 39, 1537.

[4] U. A. Argikar, J. L. Dumouchel, V. M. Kramlinger, A. L. Cirello, M. Gunduz, C. E. Dunne, B. Sohal, J. Pharm. Sci. 2017, 106, 2276.

[5] T. Zhang, C. D. Xiang, D. Gale, S. Carreiro, E. Y. Wu, E. Y. Zhang, Drug Metab. Dispos. 2008, 36, 1300.

[6] C. Kölln, S. Reichl, J. Ocul. Pharmacol. Ther. 2012, 28, 271.

[7] S. Rönkkö, K.-S. Vellonen, K. Järvinen, E. Toropainen, A. Urtti, Drug Deliv. Transl. Res. 2016, 6, 660.

[8] X. Larrea, C. De Courten, V. Feingold, J. Burger, P. Büchler, Optom. Vis. Sci. 2007, 84, 1074.

[9] M. Hornof, E. Toropainen, A. Urtti, Eur. J. Pharm. Biopharm. 2005, 60, 207.

[10] S. Reichl, J. Pharm. Pharmacol. 2008, 60, 299.

[11] D. Huh, B. D. Matthews, A. Mammoto, M. Montoya-Zavala, H. Y. Hsin, D. E. Ingber, Science (80-.). 2010, 328, 1662.

[12] J. Seo, D. Huh, in 18th Int. Conf. Miniaturized Syst. Chem. Life Sci. MicroTAS 2014, 2014.

[13] D. Bennet, Z. Estlack, T. Reid, J. Kim, Lab Chip 2018, 18, 1539.

[14] S. N. Bhatia, D. E. Ingber, Nat. Biotechnol. 2014, 32, 760.

[15] R. Abdalkader, K.-I. Kamei, Lab Chip 2020, 20, 1410.

[16] C. W. McAleer, A. Pointon, C. J. Long, R. L. Brighton, B. D. Wilkin, L. R. Bridges, N. Narasimhan Sriram, K. Fabre, R. McDougall, V. P. Muse, J. T. Mettetal, A. Srivastava, D. Williams, M. T. Schnepper, J. L. Roles, M. L. Shuler, J. J. Hickman, L. Ewart, Sci. Rep. 2019, 9, 9619.

[17] D. Bovard, A. Sandoz, K. Luettich, S. Frentzel, A. Iskandar, D. Marescotti, K. Trivedi, E. Guedj, Q. Dutertre, M. C. Peitsch, J. Hoeng, Lab Chip 2018, 18, 3814.

[18] K.-J. Jang, M. A. Otieno, J. Ronxhi, H.-K. Lim, L. Ewart, K. R. Kodella, D. B. Petropolis, G. Kulkarni, J. E. Rubins, D. Conegliano, J. Nawroth, D. Simic, W. Lam, M. Singer, E. Barale, B. Singh, M. Sonee, A. J. Streeter, C. Manthey, B. Jones, A. Srivastava, L. C. Andersson, D. Williams, H. Park, R. Barrile, J. Sliz, A. Herland, S. Haney, K. Karalis, D. E. Ingber, G. A. Hamilton, Sci. Transl. Med. 2019, 11, eaax5516.

[19] G. A. Theodoridis, H. G. Gika, I. D. Wilson, in Metabolomics Pract., Wiley-VCH Verlag GmbH & Co. KGaA, Weinheim, Germany, 2013, pp. 93–115.

[20] D. C. Sévin, T. Fuhrer, N. Zamboni, U. Sauer, Nat. Methods 2017, 14, 187.

[21] R. Abdalkader, R. Chaleckis, I. Meister, P. Zhang, C. E. Weelock, K. Kamei, Anal. Sci. 2020, advpub, DOI 10.2116/analsci.20N032.

[22] K. Kamei, Y. Koyama, Y. Tokunaga, Y. Mashimo, M. Yoshioka, C. Fockenberg, R. Mosbergen, O. Korn, C. Wells, Y. Chen, Adv. Healthc. Mater. 2016, 5, 2951.

[23] C. Kölln, S. Reichl, J. Ocul. Pharmacol. Ther. 2012, 28, 271.

[24] U. Becker, C. Ehrhardt, N. Daum, C. Baldes, U. F. Schaefer, K. W. Ruprecht, K.-J. Kim, C.-M. Lehr, J. Ocul. Pharmacol. Ther. 2007, 23, 172.

[25] M. G. Doane, A. D. Jensen, C. H. Dohlman, Am. J. Ophthalmol. 1978, 85, 383.

[26] S. P. Sugrue, J. D. Zieske, Exp. Eye Res. 1997, 64, 11.

[27] H. Yamaguchi, T. Takezawa, Drug Metab. Dispos. 2018, 46, 1684.

[28] S. P. H. Alexander, E. Kelly, N. Marrion, J. A. Peters, H. E. Benson, E. Faccenda, A. J. Pawson, J. L. Sharman, C. Southan, J. A. Davies, Br. J. Pharmacol. 2015, 172, 6110.

[29] D. C. Rees, E. Johnson, O. Lewinson, Nat. Rev. Mol. Cell Biol. 2009, 10, 218.

[30] K.-S. Vellonen, E. Mannermaa, H. Turner, M. Häkli, J. Mario Wolosin, T. Tervo, P. Honkakoski, A. Urtti, J. Pharm. Sci. 2010, 99, 1087.

[31] J.-T. Zhang, Cell Res. 2007, 17, 311.

[32] L. W. Sumner, A. Amberg, D. Barrett, M. H. Beale, R. Beger, C. A. Daykin, T. W. M. Fan, O. Fiehn, R. Goodacre, J. L. Griffin, T. Hankemeier, N. Hardy, J. Harnly, R. Higashi, J. Kopka, A. N. Lane, J. C. Lindon, P. Marriott, A. W. Nicholls, M. D. Reily, J. J. Thaden, M. R. Viant, Metabolomics 2007, 3, 211.

[33] J. Chong, O. Soufan, C. Li, I. Caraus, S. Li, G. Bourque, D. S. Wishart, J. Xia, Nucleic Acids Res. 2018, 46, W486.

[34] A. Fabregat, K. Sidiropoulos, G. Viteri, P. Marin-Garcia, P. Ping, L. Stein, P. D’Eustachio, H. Hermjakob, Bioinformatics 2018, 34, 1208.

[35] S. K. Nigam, K. T. Bush, G. Martovetsky, S.-Y. Ahn, H. C. Liu, E. Richard, V. Bhatnagar, W. Wu, Physiol. Rev. 2015, 95, 83.

[36] L. Lin, S. W. Yee, R. B. Kim, K. M. Giacomini, Nat. Rev. Drug Discov. 2015, 14, 543.

[37] E. Ganea, J. J. Harding, Curr. Eye Res. 2006, 31, 1.

[38] H. J. Gukasyan, K.-J. Kim, V. H. L. Lee, R. Kannan, Ocul. Surf. 2007, 5, 269.

[39] Y. Chen, G. Mehta, V. Vasiliou, Ocul. Surf. 2009, 7, 176.

[40] S. P. C. Cole, R. G. Deeley, Trends Pharmacol. Sci. 2006, 27, 438.

[41] H. J. Gukasyan, V. H. L. Lee, K.-J. Kim, R. Kannan, Invest. Ophthalmol. Vis. Sci. 2002, 43, 1154.

[42] T. Minich, J. Riemer, J. B. Schulz, P. Wielinga, J. Wijnholds, R. Dringen, J. Neurochem. 2006, 97, 373.

[43] J. Bai, L. Lai, H. C. Yeo, B. C. Goh, T. M. C. Tan, Int. J. Biochem. Cell Biol. 2004, 36, 247.

[44] P. Borst, C. de Wolf, K. van de Wetering, Pflügers Arch. - Eur. J. Physiol. 2007, 453, 661.

[45] A. So, B. Thorens, J. Clin. Invest. 2010, 120, 1791.

[46] K. Jäger, U. Bönisch, M. Risch, D. Worlitzsch, F. Paulsen, Investig. Opthalmology Vis. Sci. 2009, 50, 1112.

[47] N. Longo, M. Frigeni, M. Pasquali, Biochim. Biophys. Acta - Mol. Cell Res. 2016, 1863, 2422.

[48] S. Xu, J. L. Flanagan, P. A. Simmons, J. Vehige, M. D. Willcox, Q. Garrett, Mol. Vis. 2010, 16, 1823.

[49] Q. Garrett, S. Xu, P. A. Simmons, J. Vehige, J. L. Flanagan, M. D. Willcox, Investig. Opthalmology Vis. Sci. 2008, 49, 4844.

[50] P. Giesbertz, J. Ecker, A. Haag, B. Spanier, H. Daniel, J. Lipid Res. 2015, 56, 2029.

[51] F. V. Ventura, L. Ijlst, J. Ruiter, R. Ofman, C. G. Costa, C. Jakobs, M. Duran, I. T. De Almeida, L. L. Bieber, R. J. A. Wanders, Eur. J. Biochem. 1998, 253, 614.

[52] F. P. J. Diecke, L. Ma, P. Iserovich, J. Fischbarg, Biochim. Biophys. Acta - Biomembr. 2007, 1768, 2043.

[53] K. K. Williams, M. A. Watsky, Curr. Eye Res. 2004, 28, 109.

[54] K. Kamei, Y. Mashimo, Y. Koyama, C. Fockenberg, M. Nakashima, M. Nakajima, J. Li, Y. Chen, Biomed. Microdevices 2015, 17, 36.

[55] C. McQuin, A. Goodman, V. Chernyshev, L. Kamentsky, B. A. Cimini, K. W. Karhohs, M. Doan, L. Ding, S. M. Rafelski, D. Thirstrup, W. Wiegraebe, S. Singh, T. Becker, J. C. Caicedo, A. E. Carpenter, PLOS Biol. 2018, 16, e2005970.

[56] S. Naz, H. Gallart-Ayala, S. N. Reinke, C. Mathon, R. Blankley, R. Chaleckis, C. E. Wheelock, Anal. Chem. 2017, 89, 7933.

[57] R. Chaleckis, S. Naz, I. Meister, C. E. Wheelock, in Clinical Metabolomics: Methods and Protocols, (Ed: M. Giera), Springer, New York, United States 2018, pp. 45–58.

[58] I. Tada, H. Tsugawa, I. Meister, P. Zhang, R. Shu, R. Katsumi, C. E. Wheelock, M. Arita, R. Chaleckis, Metabolites 2019, 9, 251.

[59] M. C. Chambers, B. Maclean, R. Burke, D. Amodei, D. L. Ruderman, S. Neumann, L. Gatto, B. Fischer, B. Pratt, J. Egertson, K. Hoff, D. Kessner, N. Tasman, N. Shulman, B. Frewen, T. A. Baker, M.-Y. Brusniak, C. Paulse, D. Creasy, L. Flashner, K. Kani, C. Moulding, S. L. Seymour, L. M. Nuwaysir, B. Lefebvre, F. Kuhlmann, J. Roark, P. Rainer, S. Detlev, T. Hemenway, A. Huhmer, J. Langridge, B. Connolly, T. Chadick, K. Holly, J. Eckels, E. W. Deutsch, R. L. Moritz, J. E. Katz, D. B. Agus, M. MacCoss, D. L. Tabb, P. Mallick, Nat. Biotechnol. 2012, 30, 918.

[60] H. Tsugawa, T. Cajka, T. Kind, Y. Ma, B. Higgins, K. Ikeda, M. Kanazawa, J. VanderGheynst, O. Fiehn, M. Arita, Nat. Methods 2015, 12, 523.

[61] I. Meister, P. Zhang, A. Sinha, C. M. Sköld, Å. M. Wheelock, T. Izumi, R. Chaleckis, C. E. Wheelock, 2020, DOI 10.26434/chemrxiv.13489158.v1.

[62] I. Tada, R. Chaleckis, H. Tsugawa, I. Meister, P. Zhang, N. Lazarinis, B. Dahlén, C. E. Wheelock, M. Arita, Anal. Chem. 2020, 92, 11310.

[63] D. Broadhurst, R. Goodacre, S. N. Reinke, J. Kuligowski, I. D. Wilson, M. R. Lewis, W. B. Dunn, Metabolomics 2018, 14, 72.

[64] K. Haug, K. Cochrane, V. C. Nainala, M. Williams, J. Chang, K. V. Jayaseelan, C. O’Donovan, Nucleic Acids Res. 2020, 48, D440.

